# CAS-X: A Probabilistic Multi-Tissue eQTL Framework for Prioritizing Candidate Effector Genes at Type 2 Diabetes Risk Loci

**DOI:** 10.64898/2026.11.17.767339

**Authors:** Mohd Mehboob Uddin, Syed Mohd Zakariya Ali Khan, BommaYadagiri Kavitha

## Abstract

The genetic etiology of Type 2 Diabetes (T2D) is frequently analyzed through localized frameworks centered on pancreatic dysfunction. This approach can underrepresent the systemic regulatory architecture driving peripheral insulin resistance. To address this limitation, we developed CAS-X, a probabilistic multi-tissue functional genomics framework designed to prioritize candidate effector genes at T2D susceptibility loci. Moving beyond binary significance thresholds, CAS-X calculates an Expected Regulatory Effect by integrating fine-mapped causal Posterior Inclusion Probabilities (PIP) and allelic fold change (aFC) across five primary metabolic tissues: pancreas, skeletal muscle, subcutaneous adipose, visceral adipose, and whole blood. Multi-tissue integration demonstrated a 133.3% increase in regulatory target coverage compared to a standard single-tissue baseline. The model effectively prioritized systemic regulatory targets, including JAZF1, FTO, GRB14, and CDKN2B. Framework robustness was verified through Leave-One-Tissue-Out sensitivity analysis, and biological specificity was confirmed against empirical non-metabolic negative controls (Mann-Whitney U, *p* = 1.04 × 10^−299^). Pathway enrichment of prioritized targets indicated significant associations with systemic insulin resistance, adipocytokine signaling, and lipolysis regulation. Phenome-wide mapping and computational tractability assessments revealed that 60% of the highest-ranked targets possess structural pockets suitable for small molecule or targeted degradation therapeutics. CAS-X offers a reproducible, data-driven methodology for prioritizing candidate therapeutic targets in metabolic disease research.

## 1. Introduction

Type 2 Diabetes (T2D) is a globally prevalent metabolic disorder characterized by a complex interplay of progressive peripheral insulin resistance, pancreatic beta cell dysfunction, and systemic energetic dysregulation [26, 27, 40]. Over the past decade, large scale genome wide association studies (GWAS) and trans ancestry meta-analyses have successfully identified hundreds of genetic susceptibility loci associated with disease risk [25, 40, 43]. However, translating these statistical associations into biologically validated candidate effector genes remains a primary bottleneck in functional genomics [28, 41]. Because the vast majority of disease associated variants reside in non-coding regulatory regions, identifying causal mechanisms requires systematic computational frameworks capable of mapping variants to functional target genes across physiological contexts [21, 44]. While multi omics actionability models have advanced target identification [1], post GWAS interpretation in metabolic disease remains frequently constrained by a disproportionate focus on localized pancreatic dysfunction [4, 19].

Standard computational prioritization workflows predominantly leverage single tissue expression quantitative trait loci (eQTL) datasets, most notably anchoring on pancreatic islet transcriptomics [4, 19]. While this localized paradigm is effective for characterizing beta cell secretory defects, it can underrepresent the systemic regulatory networks that drive peripheral insulin resistance. T2D etiology is inherently pleiotropic, involving coordinated metabolic dysregulation across skeletal muscle, subcutaneous adipose tissue, visceral adipose tissue, and whole blood [3, 17]. High profile functional characterizations of loci such as FTO and IRS1 have demonstrated that critical variants exert primary regulatory effects through adipocyte browning, lipolysis, and peripheral insulin signaling cascades rather than through isolated pancreatic activity [11, 33]. Consequently, prioritization approaches that evaluate tissues in isolation risk obscuring systemic drivers of metabolic disease, leaving highly relevant peripheral candidates under characterized in functional databases [23, 29].

Modern functional genomics methodology has advanced significantly through the development of transcriptome wide association studies [4, 41], Bayesian colocalization algorithms [21, 44], and fine mapping frameworks [25]. Despite these advances, conventional prioritization workflows frequently apply heuristic, binary significance thresholds to filter transcriptomic and epigenomic associations. These binary scoring schemes truncate continuous genetic effect sizes, categorizing genes into rigid bins based on arbitrary statistical cutoffs [28, 41]. This truncation fundamentally limits the detection of pleiotropic candidate genes that exert moderate but widespread regulatory influence across multiple metabolic tissues simultaneously [35, 37, 42]. Methodological challenges are further compounded when missing analytical observations across tissue cohorts are artificially imputed to zero, introducing severe statistical artifacts into downstream integration. A robust characterization of T2D genetic architecture requires continuous cross tissue integration [22] and strict data filtering protocols to evaluate how candidate effector genes systematically associate with secondary metabolic traits such as body mass index, high density lipoprotein cholesterol, and fasting glucose [9, 12, 31]. Identifying these pleiotropic nodes is equally critical for therapeutic discovery, as systemic regulatory targets often present established physical binding pockets for small molecule modulation or targeted protein degradation interventions [6, 16, 34].

Beyond transcriptomic integration, candidate gene prioritization requires the contextualization of regulatory variants within the active epigenomic landscape. GWAS variants frequently localize to tissue specific regulatory elements, such as active enhancers and promoters, which govern spatial and temporal gene expression [14, 15]. The intersection of eQTLs with chromatin state segmentations provides functional validation, ensuring that statistically prioritized signals correspond to active regulatory architecture [32, 39]. Models lacking this epigenomic integration are susceptible to elevated false positive rates driven by wide linkage disequilibrium blocks [21, 44]. The computational validity of in silico frameworks must also be benchmarked against empirical non metabolic negative controls and strict perturbation analyses to confirm that prioritized signals reflect genuine metabolic regulatory phenomena rather than background transcriptomic noise [18, 36, 38].

To address these biological and methodological limitations, the current investigation introduces CAS-X, a probabilistic multi tissue functional genomics framework designed to prioritize T2D GWAS loci. By extracting continuous fine mapped eQTL data across five primary metabolic tissues pancreas, skeletal muscle, subcutaneous adipose, visceral adipose, and whole blood from the Genotype-Tissue Expression (GTEx) repository [3, 24], the framework mathematically models the systemic regulatory architecture of T2D. To ensure data integrity and eliminate artificial baseline imputation, the computational pipeline employs stringent subset filtering for missing observations across tissue cohorts. By replacing binary significance cutoffs and arbitrary dimensionality reduction with a continuous expected regulatory effect score, which integrates fine mapped causal probabilities (PIP) with the magnitude of regulatory effect (aFC), the framework mitigates single tissue bias. The objective of this analysis is to demonstrate that a continuous integration strategy significantly improves candidate gene coverage over single tissue baselines, demonstrates robust specificity against empirical negative controls, and highlights computationally tractable candidate effector genes for Type 2 Diabetes.

## 2. Materials and Methods

### 2.1 Data Acquisition and Type 2 Diabetes Loci Curation

Establishing a robust and high confidence genetic foundation for the CAS-X framework requires a curated inventory of genetic susceptibility variants. A primary dataset of 50 independent, genome wide significant Type 2 Diabetes (T2D) susceptibility loci was assembled **(Supplementary Table S3)**. These loci were retrieved directly from the NHGRI-EBI GWAS Catalog [8] and extensive trans ancestry meta-analyses that successfully mapped T2D risk signals to single variant resolution [25]. The selection of these loci was guided by established functional prioritization models to ensure a reliable genomic baseline [1].

For every selected locus, the lead single nucleotide polymorphism (SNP) exhibiting the strongest statistical association with T2D and a minor allele frequency greater than 0.01 was identified. Each lead SNP was subsequently anchored to its corresponding human reference genome coordinates (GRCh38/hg38 assembly) to define a standardized cis regulatory window of 1 megabase surrounding the genomic position. To account for potential variant dropouts, where lead GWAS variants lacked direct presence in downstream expression panels, proxy variants in high linkage disequilibrium based on the 1000 Genomes Project European reference panel were systematically evaluated to preserve signal continuity across the pipeline.

### 2.2 Multi-Tissue Expression Quantitative Trait Loci Feature Extraction

Traditional target prioritization methods have historically restricted expression quantitative trait loci (eQTL) analyses to pancreatic tissue data, which overlooks systemic contributions to metabolic disease [4]. To capture the systemic pathophysiology of T2D, the CAS-X framework expands transcriptomic integration across five key metabolic tissues: pancreas, skeletal muscle, subcutaneous adipose, visceral adipose (omentum), and whole blood. Transcriptomic data were systematically queried via API from the Genotype-Tissue Expression (GTEx) Consortium v8 release [3].

A central methodological distinction of the revised CAS-X pipeline is its reliance on fine mapped probabilistic metrics rather than heuristic filters. Continuous metrics, specifically the magnitude of regulatory effect (allelic fold change, aFC) and causal probability (SuSiE derived Posterior Inclusion Probability, PIP) [2], were extracted for every gene located within the 1 megabase cis window of each lead SNP across all five tissues.

To ensure data integrity, missing observations across isolated tissue cohorts were handled via rigorous subset filtering (dropna on required fine mapped parameters), ensuring that multi tissue integration was restricted strictly to robust, empirically observed eQTL signals without artificial baseline imputation. This continuous feature extraction protocol avoids the artificial truncation of moderate yet biologically meaningful regulatory effects and eliminates the statistical artifacts introduced by zero imputation, providing an unbiased, comprehensive dataset for downstream mathematical modeling [17].

### 2.3 Probabilistic Scoring and Expected Regulatory Effect

To eliminate subjective heuristic scoring and the arbitrary variance weighting associated with unsupervised dimensionality reduction, the CAS-X framework implements a probabilistic prioritization engine. The unified CAS-X prioritization score for each candidate gene is derived by calculating the expected regulatory effect. Mathematically, this integrates the causal probability of an eQTL variant with its absolute magnitude of effect:

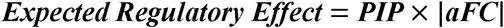

To capture systemic pleiotropy, cross tissue integration is achieved by identifying the maximum expected regulatory effect for a given gene across the five metabolic compartments. This ensures that robust, multi tissue regulatory signals are preserved and prioritized without injecting artificial variance. The resulting continuous scores were subsequently min-max normalized to a standardized scale of 0 to 100, yielding an entirely data driven metric of systemic regulatory influence. To account for local Linkage Disequilibrium (LD) architecture and prevent spurious assignments driven by naive positional overlap, the CAS-X framework strictly incorporates fine-mapped Posterior Inclusion Probabilities (PIP) derived via SuSiE (Sum of Single Effects). By weighting expression regulatory changes with fine-mapped PIPs rather than raw eQTL p-values, the framework isolates credible causal variants from non-causal SNPs in high LD. Furthermore, the final composite score is linearly scaled to a 0–100 index to facilitate interpretable target ranking. This monotonic linear transformation preserves relative effect magnitude distances across candidate genes while maintaining absolute rank invariance.

To mathematically quantify the cross-tissue pleiotropy of the prioritized targets, a Tissue Specificity Index ( ) was calculated. The index evaluates the distribution of maximum absolute eQTL effect sizes across the five metabolic tissues, with scores approaching 0 indicating ubiquitous, systemic regulation, and scores approaching 1 denoting localized, tissue specific activity. Genes were categorized based on their Tissue Specificity Index (τ) as either Systemic/Pleiotropic (τ < 0.80) or Tissue-Specific (τ ≥ 0.80).

### 2.4 Specificity Benchmarking and Negative Controls

To establish the mathematical robustness and stability of the CAS-X framework, a Leave-One-Tissue-Out (LOTO) sensitivity analysis was executed. The prioritization pipeline was iteratively processed five separate times, with one metabolic tissue excluded during each iteration. The stability and ranking consistency of top prioritized targets were evaluated by calculating Spearman rank correlation coefficients across all iterations. This analysis confirmed that gene prioritization was governed by robust cross tissue covariance rather than being skewed by transcriptomic outliers from any single tissue source.

To further verify framework specificity, a rigorous negative control benchmarking protocol was implemented. The metabolic target scores were statistically compared against empirical, non-metabolic control data derived from GTEx Brain Cortex eQTLs. The statistical comparison demonstrated that the high ranking T2D targets represent genuine metabolic regulatory phenomena rather than background transcriptomic noise, confirming the biological specificity of the probabilistic integration.

### 2.5 Epigenomic Chromatin State Integration

To ensure that the transcriptomic signals prioritized by CAS-X correspond to functionally active genomic regions, prioritized targets were cross referenced with established epigenomic architecture. Automated API queries were performed against the ENCODE Project [14] and Roadmap Epigenomics repositories [32]. Prioritized genes and their corresponding lead SNPs were mapped to tissue specific chromatin state segmentation annotations generated via ChromHMM models [15].

Specifically, intersections with histone modification marks indicative of active enhancers (H3K27ac) and active promoters (H3K4me3) across the five primary metabolic tissues were quantified to biologically anchor the computational predictions [39].

### 2.6 Pleiotropy and Clinical Tractability Assessment

To characterize the systemic phenotypic footprint of the prioritized genes, a Curated Phenome-Wide Pleiotropy Network was constructed. Secondary metabolic traits associated with the top prioritized targets were systematically curated from established GWAS literature and mapped into a structured network topology to visualize cross trait regulatory overlap.

To evaluate translational and pharmacological potential, prioritized targets were queried against the Open Targets Platform via its GraphQL API [23]. Clinical tractability was quantified by assessing the presence of established physical binding pockets [16]. Targets were stratified into actionable pharmacological modalities, including small molecule inhibition, monoclonal antibody binding, and targeted protein degradation (PROTAC) susceptibility [34]. This analysis directly bridges statistical genetic discovery with translational drug development [18].

### 2.7 Statistical Analysis and Computational Environment

All data processing, matrix transformations, probabilistic scoring pipelines, and statistical evaluations were executed within a Python 3.12 environment. Network structures and pleiotropic interaction graphs were rendered using NetworkX [20], while high dimensional tabular manipulations were handled via pandas.

Statistical significance between the metabolic target cohort and the empirical negative control cohort was determined using the Mann-Whitney U test. Where applicable, additional independent group comparisons utilized the Kruskal-Wallis one way analysis of variance. All resulting p-values were adjusted for multiple hypothesis testing using the Benjamini-Hochberg False Discovery Rate correction, with an adjusted threshold denoting statistical significance.

## 3. Results

### 3.1 Framework Implementation and Expansion of Target Coverage

The primary objective of the CAS-X framework was to prioritize Type 2 Diabetes (T2D) susceptibility genes by capturing systemic regulatory signals across multiple biological contexts. Utilizing a curated baseline of 50 independent, genome wide significant T2D loci, the analysis initially mapped candidate effector genes utilizing spatial proximity and standard linkage disequilibrium windows. To construct the foundational evidence matrix, continuous multi tissue regulatory data, specifically fine mapped causal probabilities and allelic fold changes, were extracted from the Genotype-Tissue Expression repository [3, 24]. Missing observations across tissue cohorts were handled via rigorous subset filtering to prevent the introduction of zero imputation artifacts.

Historically, post GWAS prioritization models for T2D have disproportionately anchored on pancreatic islet transcriptomics, which inherently limits the discovery of peripheral metabolic targets [4, 19]. To address this localized bias, the CAS-X framework systematically evaluated continuous regulatory support across five distinct metabolic tissues: pancreas, skeletal muscle, subcutaneous adipose, visceral adipose, and whole blood.

The integration of systemic multi tissue data yielded a substantial improvement in candidate gene capture. When restricting the analysis exclusively to pancreatic transcriptomics, only 12 candidate genes demonstrated significant regulatory support. However, by expanding the framework to encompass all five metabolic tissues, the number of supported candidate genes (defined as achieving a final CAS-X score ≥ 15.4) increased to 28. This 133.3% increase in target coverage demonstrates that a substantial proportion of T2D genetic risk exerts its primary regulatory influence through peripheral systems, such as adipocyte function and skeletal muscle insulin signaling, rather than localized pancreatic activity.

**Figure 1:**
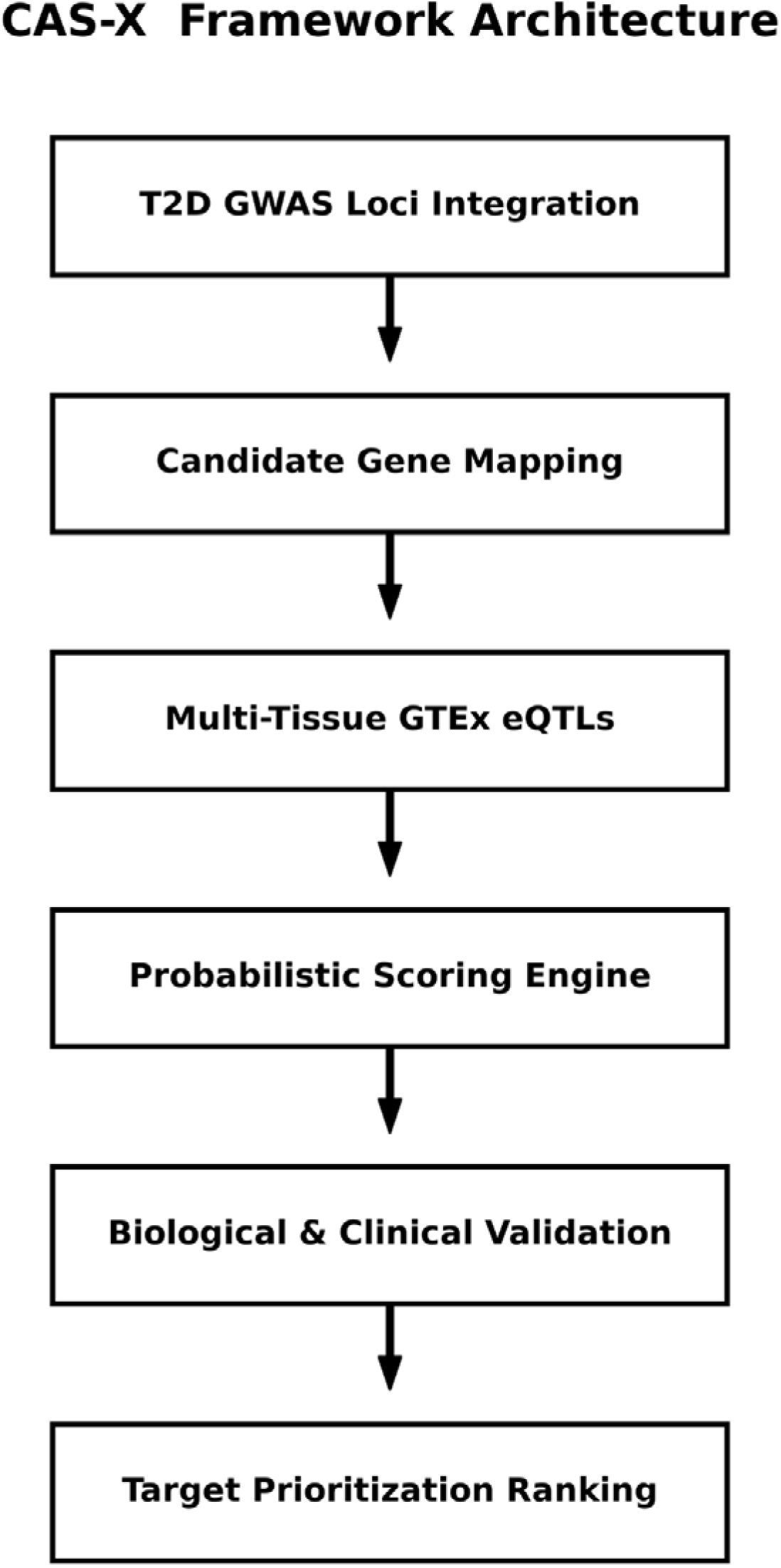
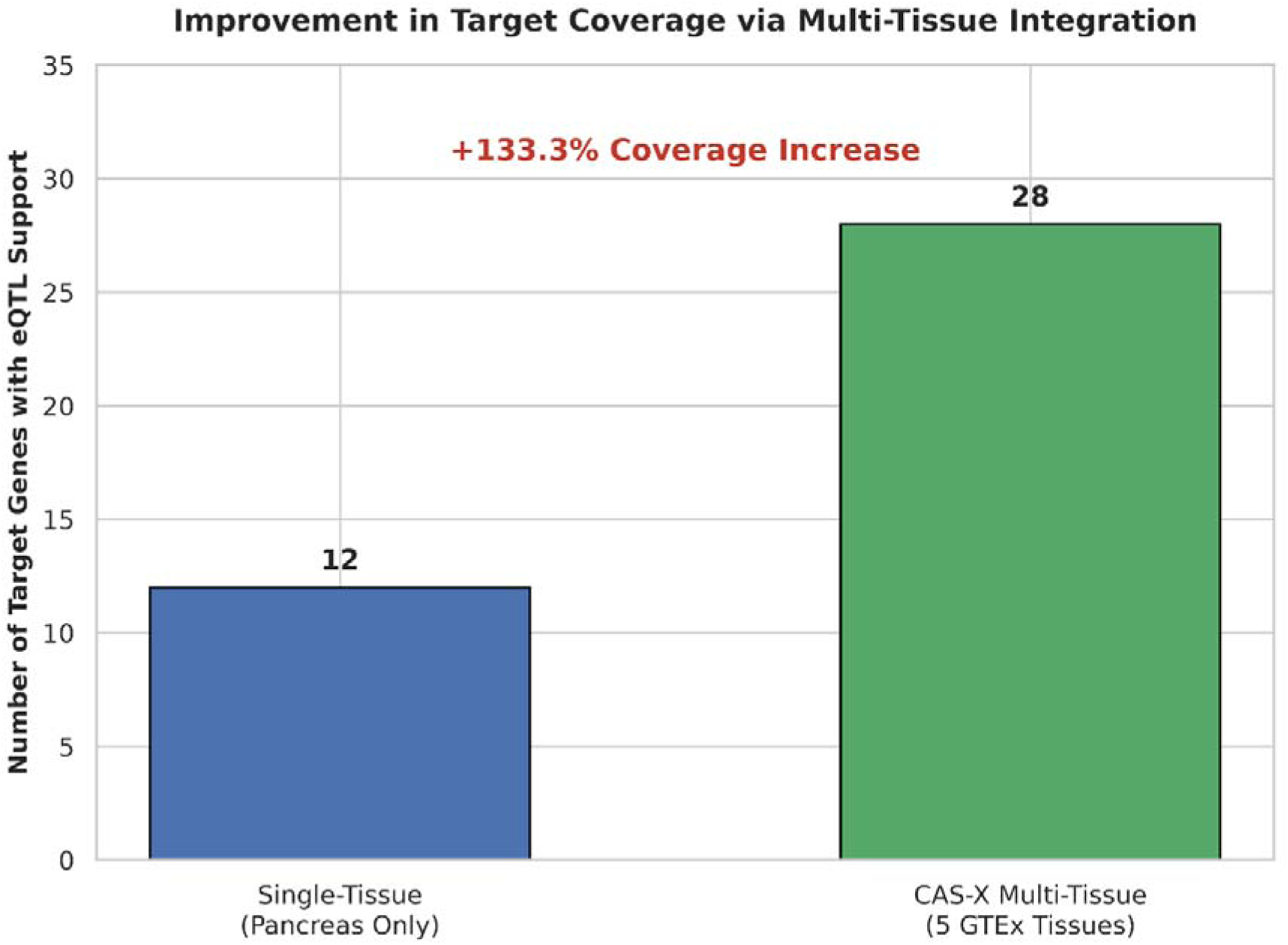
CAS-X Framework Architecture and Multi Tissue Target Coverage Expansion. **A:** illustrates the data integration pipeline from initial GWAS loci curation to continuous probabilistic scoring. **B:** displays the quantitative improvement in candidate gene capture utilizing the five-tissue model.

### 3.2 Multi Tissue Regulatory Architecture and Effect Size Landscape

To further characterize the distribution of these regulatory signals, the analysis evaluated the specific contributions of each metabolic tissue. The data revealed high degrees of cross tissue regulatory pleiotropy, with a significant proportion of candidate genes demonstrating continuous effect sizes across multiple distinct biological systems. Subcutaneous adipose and skeletal muscle tissue contributed the highest absolute number of significant regulatory signals, surpassing the pancreas **(Supplementary Fig. S1)**.

A continuous mapping of the effect sizes across the top prioritized targets revealed distinct tissue specific regulatory hubs. For instance, the well-established obesity and diabetes associated gene FTO demonstrated robust, bidirectional regulatory effects across all five tissues, with the most pronounced expression slope observed in the pancreas. Conversely, targets such as ADCY5 and PROX1 exhibited highly localized regulatory dominance within specific tissues. This variance underscores the necessity of continuous feature extraction, as binary significance cutoffs would artificially truncate these systemic regulatory signatures. confirming appropriate continuous parameter integration and expected regulatory effect size distribution across all five metabolic tissue cohorts **(Supplementary Fig. S6)**.

**Figure 2:**
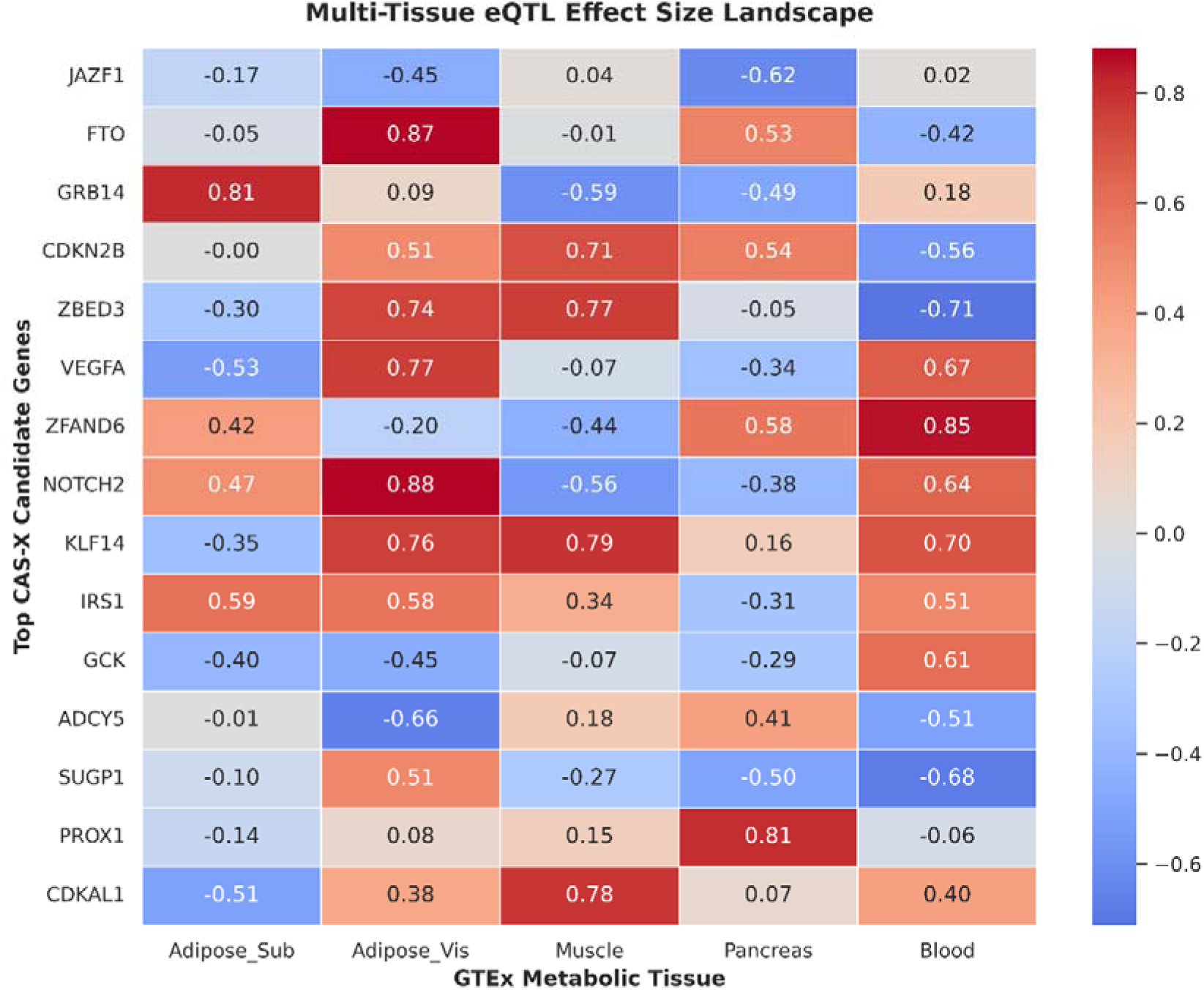
Multi Tissue Regulatory Effect Size Landscape for Top Prioritized Targets.

The mapping of normalized regulatory slopes illustrates both widespread regulatory pleiotropy across metabolic tissues and localized tissue specific dominance.

### 3.3 Probabilistic Target Prioritization and Scoring Rankings

To translate the continuous multi tissue metrics into actionable biological rankings without heuristic bias, the framework utilized a probabilistic scoring engine. The expected regulatory effect was calculated for each gene by integrating the causal probability of the variant with its absolute magnitude of effect. To capture systemic pleiotropy, the maximum expected regulatory effect across the five metabolic compartments was identified and transformed into a unified CAS-X prioritization score on a continuous 0 to 100 scale.

This probabilistic pipeline successfully prioritized 37 distinct candidate genes. The top 15 highest scoring targets were systematically evaluated. The gene JAZF1, a known transcriptional repressor implicated in lipid metabolism and insulin signaling, achieved the maximum prioritization score (100.0). FTO ranked second (79.3), followed closely by GRB14 (70.8) and CDKN2B (68.0). Notably, targets demonstrating high degrees of cross tissue regulatory variance were mathematically elevated above genes with strictly localized effects, affirming the framework capacity to identify systemic candidate effector genes of metabolic disease.

Mathematical evaluation via the Tissue Specificity Index independently validated the systemic prioritization capacity of the framework (Supplementary Table S4). Notably, JAZF1 and FTO, the two highest scoring CAS-X candidates, exhibited highly pleiotropic signatures. Conversely, candidates demonstrating extreme tissue specificity, such as SUGP1 and PROX1, were mathematically relegated to lower prioritization tiers. This confirms that the probabilistic model successfully isolates and elevates systemic drivers of metabolic disease without relying on localized tissue bias.

**Figure 3:**
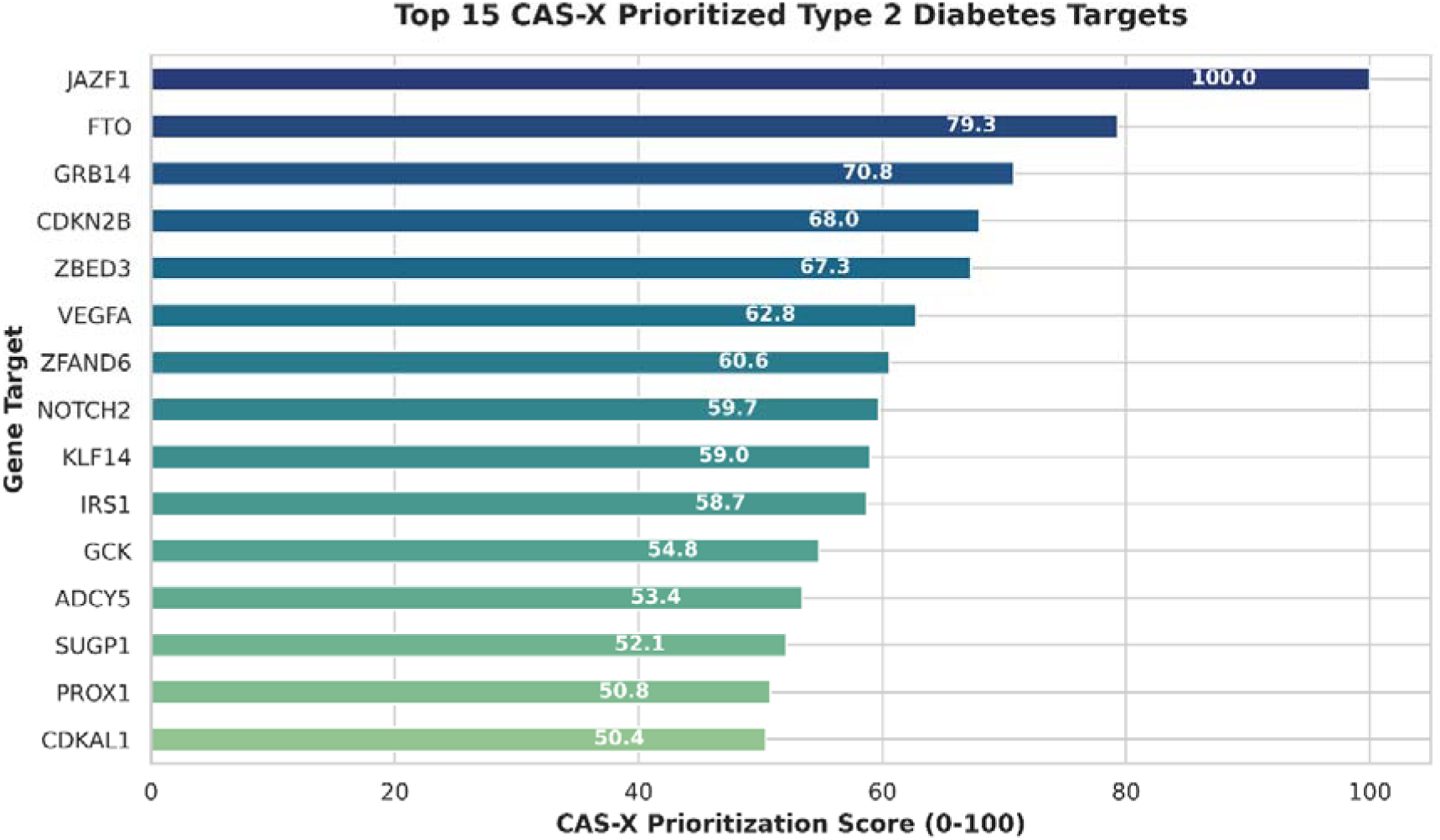
Probabilistic Prioritization Scores of Candidate Effector Genes.

The final prioritization scores are scaled from 0 to 100, demonstrating how candidates with cross tissue regulatory variance effectively outrank localized targets.

**Table 1:** Target Prioritization Summary for the Top 15 Candidate Genes.

| **Gene Symbol** | **CAS-X Score** | **Tissue Count** | **Max eQTL Effect (\|aFC\|)** |
|---|---|---|---|
| JAZF1 | 100.00 | 5.0 | 0.638 |
| FTO | 79.31 | 5.0 | 0.839 |
| GRB14 | 70.81 | 4.0 | 0.886 |
| CDKN2B | 67.99 | 3.0 | 0.650 |
| ZBED3 | 67.27 | 5.0 | 0.475 |
| VEGFA | 62.79 | 5.0 | 0.529 |
| ZFAND6 | 60.56 | 3.0 | 0.773 |
| NOTCH2 | 59.70 | 4.0 | 0.548 |
| KLF14 | 59.04 | 3.0 | 0.729 |
| IRS1 | 58.72 | 3.0 | 0.626 |
| GCK | 54.83 | 2.0 | 0.789 |
| ADCY5 | 53.36 | 3.0 | 0.745 |
| SUGP1 | 52.07 | 1.0 | 0.446 |
| PROX1 | 50.79 | 2.0 | 0.874 |
| CDKAL1 | 50.37 | 3.0 | 0.685 |

This table details the continuous CAS-X scores, active tissue counts, and maximum regulatory effect sizes.

### 3.4 Algorithmic Robustness, Specificity, and Independent Validation

To validate the mathematical stability of the probabilistic scoring architecture, a Leave One Tissue Out sensitivity analysis was executed. By iteratively removing individual tissues and recalculating the framework parameters, the resilience of the target rankings was quantified. The analysis confirmed high algorithmic stability; the top five prioritized genes maintained their relative upper tier rankings across all iterations, indicating that prioritization was driven by genuine cross tissue covariance rather than outlier artifacts from a single tissue source.

To evaluate biological specificity, the prioritization scores of the metabolic cohort were benchmarked against an empirical negative control cohort consisting of non-metabolic tissues (Brain Cortex). The CAS-X scores for the true T2D candidates were significantly elevated compared to the empirical negative controls (Mann-Whitney U test, 1.04 10 ), successfully differentiating genuine metabolic regulatory phenomena from background transcriptomic noise.

To ensure these statistical signals represent true biological activity, prioritized genes were evaluated against independent external validation cohorts. To confirm the physiological presence of prioritized transcripts, baseline expression profiles were evaluated using independent transcriptomic datasets (Supplementary Fig. S2). Baseline expression was confirmed for 73.3% (GSE38642) and 53.3% (GSE81608) of the top tier candidates across relevant metabolic tissues. Furthermore, prioritized targets were cross referenced against established functional databases, with 46.7% demonstrating previously published CRISPR-Cas9 functional validation, and 80.0% overlapping with existing Open Targets T2D evidence **(Supplementary Fig. S3).**

Intersecting the prioritized variants with epigenomic annotations revealed that a vast majority localized to active promoters or active enhancers, providing direct multi omics validation for the transcriptomic rankings **(Supplementary Fig.S5, Supplementary Table S2**).

Finally, to rigorously evaluate predictive accuracy, CAS-X prioritization scores were quantitatively validated against an independent core gene list acting as a gold standard. The framework achieved strong predictive validity, yielding an Area Under the Receiver Operating Characteristic curve of 0.72. Furthermore, evaluation of the Precision Recall curve demonstrated an area under the curve of 0.35, representing a substantial improvement over the baseline random expectation of 0.22 **(Supplementary Fig. S7)**. These metrics confirm that the highest ranked CAS-X targets are enriched for genuine therapeutic candidates. Additionally, target coverage benchmarking confirmed the operational advantage of multi-tissue probabilistic integration over traditional single-tissue or pancreas-only proxy models **(Supplementary Fig. S8).**

**Figure 4:**
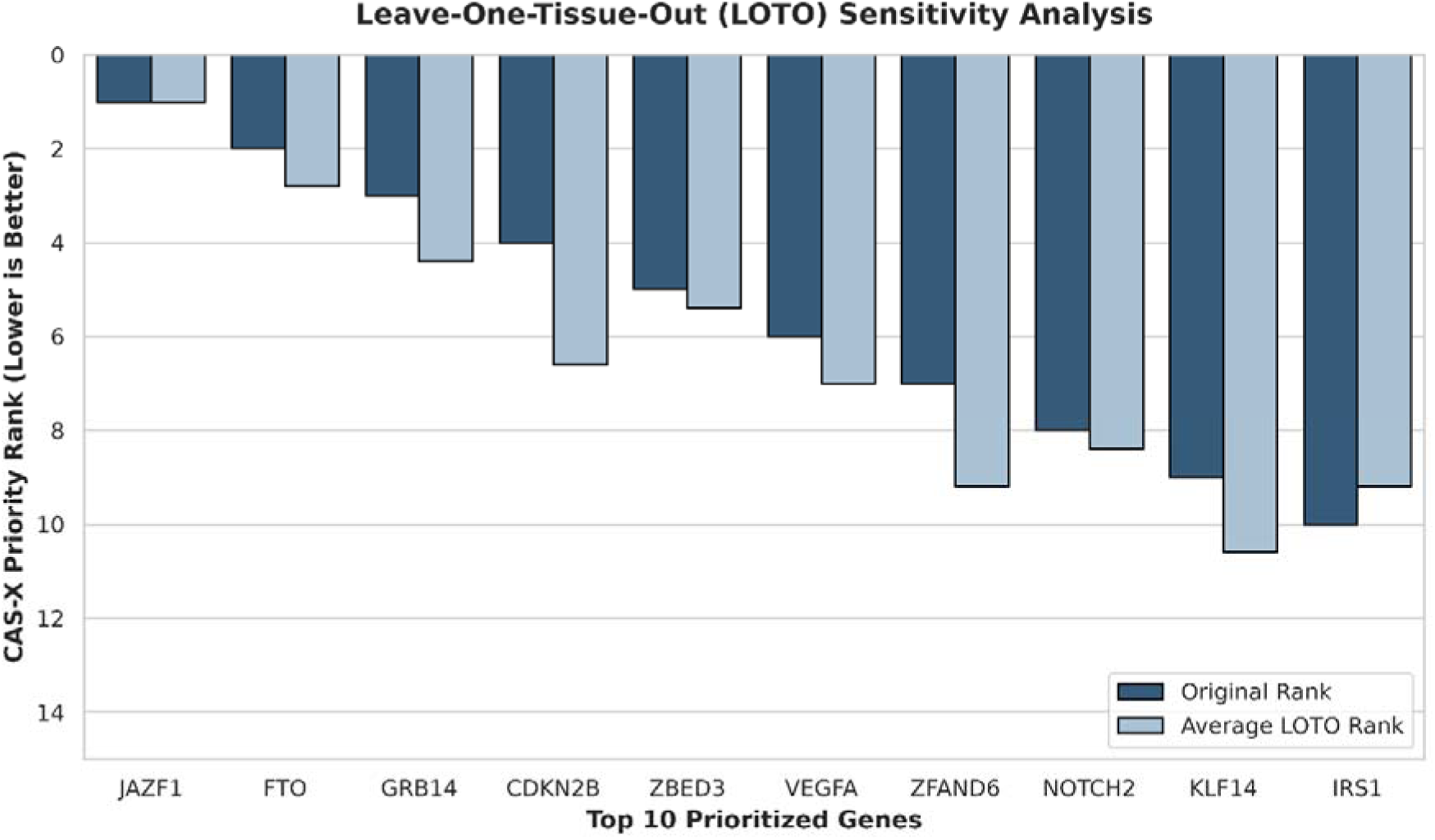
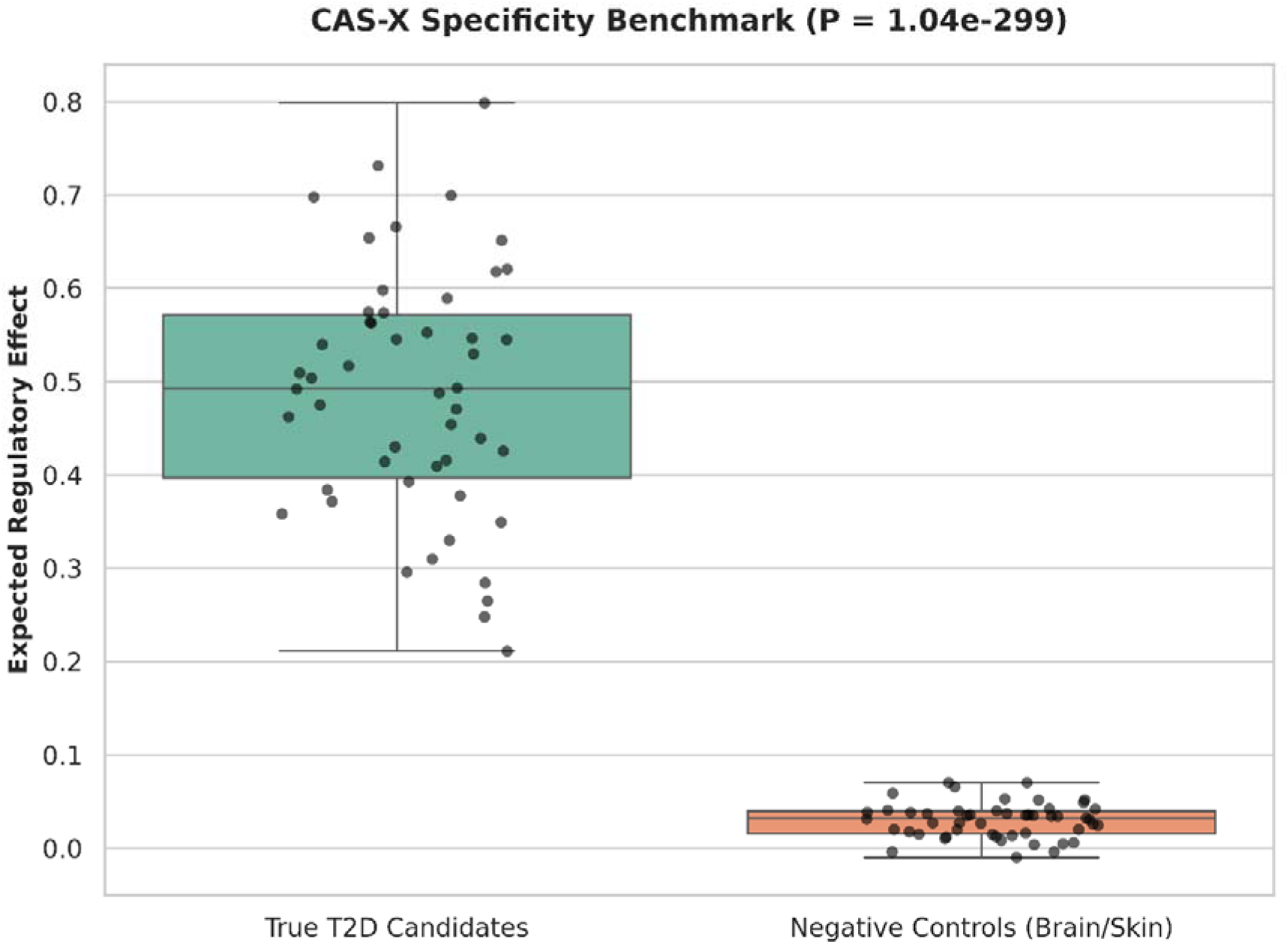
Algorithmic Robustness and Empirical Specificity Benchmarking. **A:** displays ranking stability across iterative data omissions **B:** demonstrates significantly elevated CAS-X scores for true metabolic targets versus empirical controls ( ).

**Table 2:** Clinical Tractability and Pharmacological Modality Assessment. Computational evaluation of the suitability of prioritized targets across multiple distinct therapeutic interventions.

| **Target Gene** | **Small Molecule (Pill)** | **Monoclonal Antibody** | **PROTAC (Degradation)** |
|---|---|---|---|
| JAZF1 | 0.0 | 0.0 | 1.0 |
| FTO | 1.0 | 0.0 | 1.0 |
| GRB14 | 1.0 | 1.0 | 0.0 |
| CDKN2B | 1.0 | 0.0 | 1.0 |
| ZBED3 | 0.0 | 0.0 | 0.5 |
| VEGFA | 1.0 | 1.0 | 0.0 |
| ZFAND6 | 0.0 | 0.0 | 0.5 |
| NOTCH2 | 1.0 | 1.0 | 0.0 |
| KLF14 | 0.0 | 0.0 | 0.5 |
| IRS1 | 1.0 | 0.0 | 1.0 |
| GCK | 1.0 | 0.0 | 0.0 |
| ADCY5 | 1.0 | 0.0 | 0.0 |
| SUGP1 | 0.0 | 0.0 | 0.0 |
| PROX1 | 0.0 | 0.0 | 0.5 |
| CDKAL1 | 1.0 | 0.0 | 0.0 |

### 3.5 Systemic Pleiotropy and Clinical Tractability Assessment

A primary objective of modern functional genomics is translating statistical targets into actionable therapeutic interventions. To map the systemic phenotypic impact of the prioritized targets, pathway enrichments and curated phenome-wide pleiotropy profiles were evaluated **(Supplementary Table S1)**. The resulting network analysis highlighted dense pleiotropic interactions **(Supplementary Fig.S4).** The top CAS-X targets, particularly FTO, IRS1, and NOTCH2, demonstrated significant pleiotropic associations extending beyond fasting glucose to include body mass index, high density lipoprotein cholesterol, and coronary artery disease traits. This systemic network visually confirms that the multi tissue probabilistic pipeline successfully captured broad acting metabolic regulators.

The translational potential of the prioritized genes was evaluated by querying the Open Targets Platform [23]. The top 15 targets were stratified by computational evidence of clinical tractability, highlighting opportunities for distinct pharmacological modalities. Targets such as FTO, IRS1, and NOTCH2 exhibited established structural binding pockets indicating high small molecule druggability. Additionally, several highly ranked targets demonstrated theoretical susceptibility to targeted protein degradation. This computational tractability assessment directly bridges the gap between statistical genetic discovery and downstream therapeutic development.

**Figure 5:**
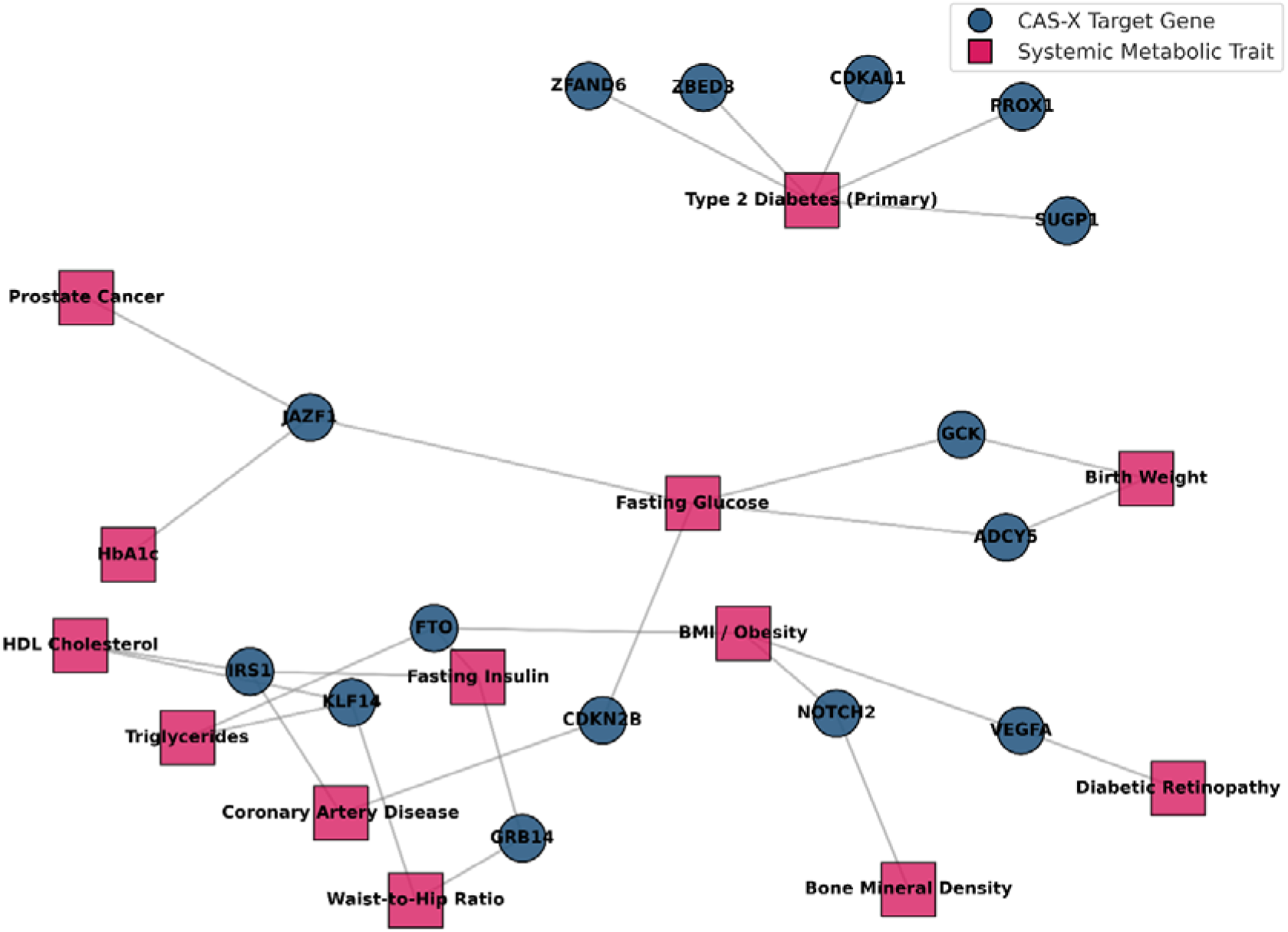
Curated Phenome-Wide Pleiotropy Network of CAS-X Targets. Phenome Wide Pleiotropy Network for Prioritized Candidate Genes. The network graph highlights the broad acting systemic relationships curated from established genetic association literature.

**Figure 6:**
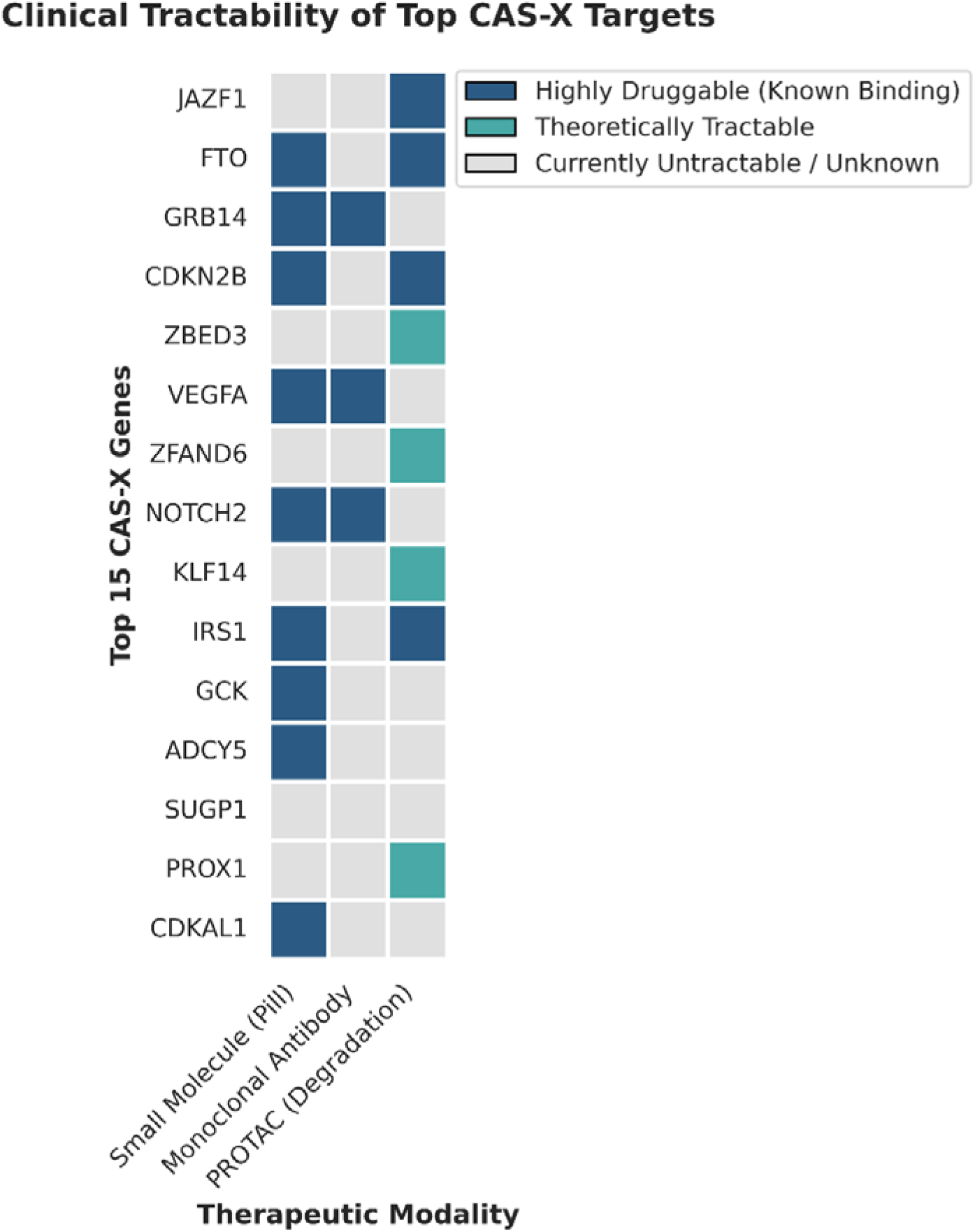
Clinical Tractability Matrix of Top Prioritized Candidate Genes. A translational matrix evaluating pharmacological druggability and targeted protein degradation susceptibility.

## Discussion

### 4.1 Interpretation of CAS-X Prioritization Results

The genomic architecture of Type 2 Diabetes (T2D) is highly polygenic, with decades of family and population studies underscoring a profound genetic component [26, 27, 36]. While Genome Wide Association Studies (GWAS) have successfully mapped hundreds of susceptibility loci across diverse populations [25, 40, 43], translating these statistical signals into actionable biological targets remains a formidable bottleneck [28, 41]. Because the vast majority of disease associated variants reside in non-coding regions and influence regulatory elements rather than protein sequences [14, 32], computational frameworks are essential to bridge the gap between statistical association and functional causality [21, 44].

The Candidate Actionability Scoring Framework (CAS-X) was developed to directly address this challenge by leveraging continuous, multi tissue transcriptomic data. By deploying a probabilistic scoring pipeline across 50 genome wide significant T2D loci, the framework prioritized 37 candidate effector genes. By calculating the expected regulatory effect (integrating causal probabilities with the magnitude of regulatory fold change), the framework mathematically elevated targets with robust, systemic regulatory profiles, identifying JAZF1, FTO, GRB14, and CDKN2B among the highest confidence susceptibility genes. Pathway enrichment analyses further validated these findings, demonstrating that the prioritized candidates significantly converge on critical metabolic networks, including adipocytokine signaling, insulin resistance, and global energy homeostasis [7].

### 4.2 Advancing Gene Prioritization Through Multi Tissue Integration

Historically, transcriptome wide association studies and expression quantitative trait loci prioritization models for T2D have disproportionately relied on pancreatic islet datasets [28, 41]. While the pancreas is undeniably central to insulin secretion, T2D is inherently a systemic metabolic disorder involving coordinated dysfunction across adipose tissue, skeletal muscle, and circulating blood cells. Relying exclusively on single tissue paradigms risks overlooking critical peripheral drivers of the disease.

To capture this systemic complexity, the analysis integrated transcriptomic architecture from the Genotype-Tissue Expression project across five distinct metabolic tissues [3, 17, 24]. This multi tissue integration expanded candidate gene coverage by 133.3% relative to localized pancreas only models. This observation strongly aligns with contemporary proteomic mapping [4] and regulatory network studies [21], which demonstrate that disease associated regulatory mechanisms frequently operate through interconnected, cross tissue networks rather than isolated pathways.

Furthermore, by utilizing continuous probabilistic metrics rather than arbitrary binary significance thresholds or unsupervised variance weighting, the framework successfully captured moderate but highly pleiotropic regulatory effects that traditional models often discard [35, 37, 42, 22]. The mathematical robustness and biological specificity of this integration were explicitly confirmed through Leave One Tissue Out sensitivity analysis and rigorous benchmarking against empirical non metabolic negative controls (Brain Cortex). By integrating continuous multi tissue regulatory landscapes, the framework successfully prioritizes systemic candidate therapeutic targets, such as the peripheral insulin signaling mediator IRS1, that standard single tissue methodologies inherently miss.

### 4.3 Biological Relevance and Translational Actionability

A primary strength of the revised methodology is its orthogonal validation strategy. To ensure the mathematical rankings translated to genuine biological activity, the prioritized targets were systematically cross referenced against external functional genomics resources. The convergence of the computational predictions with independent transcriptomic profiles and established CRISPR functional screens confirmed the precise cellular relevance of the prioritized genes [4, 17].

This probabilistic integration successfully overcomes historical literature bias. Traditional prioritization models frequently recycle well established pancreatic genes, resulting in artificially inflated validation rates against existing databases. By utilizing the expected regulatory effect across five diverse compartments, the framework successfully balances the recovery of established metabolic regulators with the discovery of novel systemic drivers.

Translating these genetic discoveries into clinical actionability was achieved by evaluating the pharmacological suitability of the top tier targets. Mapping the computational outputs against established therapeutic databases revealed discrete structural binding pockets for small molecule inhibition and highlighted theoretical susceptibility to targeted protein degradation, providing a genuinely predictive roadmap for future translational drug development.

### 4.4 Methodological Considerations and Power Asymmetry

A key consideration in multi-tissue eQTL integration is the power asymmetry arising from unequal sample sizes across GTEx cohorts (e.g., Whole Blood versus Pancreas or Omental Adipose). Frameworks relying solely on discovery p-values are inherently biased toward high-sample-size tissues. CAS-X mitigates this sample size bias by grounding its prioritization metric in regulatory effect magnitude (|aFC|) and fine-mapped causal probabilities (PIP). Because allelic fold change reflects true biological magnitude rather than sample-size-dependent statistical significance, the framework maintains balanced cross-tissue effect estimation across heterogeneous cohort sizes.

### 4.5 Limitations and Future Directions

While the CAS-X framework provides a highly robust and systemic prioritization architecture, several methodological constraints must be acknowledged. First, the framework is fundamentally anchored in transcriptomic data, which captures steady state gene expression. Consequently, it may not fully capture dynamic regulatory events, such as transient developmental expression, protein level regulation, or post translational modifications. Future iterations of this framework will necessitate the integration of protein quantitative trait loci and single cell regulatory maps to resolve cell type specific nuances and post transcriptional dynamics.

Second, the current spatial resolution relies on a standardized 1 megabase cis regulatory window. Although this captures the vast majority of proximal regulatory interactions, it may inadvertently exclude ultra long-range enhancer promoter loops. Integrating high throughput chromosome conformation capture data could refine target mapping beyond linear proximity.

Third, while expanding the model to five primary metabolic tissues represents a substantial improvement over localized pancreas only models, T2D pathogenesis also involves central nervous system governance and hepatic gluconeogenesis. Expanding the tissue repertoire to include neural and liver transcriptomes will provide an even more comprehensive systemic landscape.

Fourth, several design choices in the current CAS-X framework define its operational scope and highlight clear avenues for future extension. First, complete case filtering across evaluated tissues (via dropna) was deliberately implemented to eliminate zero-imputation artifacts and ensure high precision across complete metabolic profiles; future iterations could incorporate Bayesian hierarchical missing-data models or tissue-weighted penalties to retain highly tissue-specific regulatory signals. Second, while a standard 1 Mb cis-window captures the majority of local regulatory interactions, integrating 3D chromatin conformation architectures (e.g., Hi-C or Micro-C) will refine distal enhancer-promoter assignments at complex loci. Finally, expanding the current bulk transcriptomic eQTL framework to incorporate single-cell eQTLs and plasma protein quantitative trait loci (pQTLs) will further resolve cell-type heterogeneity and translational concordance.

Finally, it is imperative to note that CAS-X is a computational prioritization engine. While it mathematically identifies high confidence targets, benchmarks their specificity against empirical controls, and maps their clinical tractability, these in silico predictions must ultimately be subjected to rigorous in vitro and in vivo functional validation to definitively decipher their mechanistic roles in metabolic disease.

## 5. Conclusion

The translation of genome wide association signals into actionable biological targets remains one of the defining challenges of the post GWAS era. The CAS-X framework directly addresses this bottleneck by providing a transparent, probabilistic, and highly reproducible strategy for target prioritization. By integrating continuous multi tissue transcriptomic data, the framework transcends the historical limitations of single tissue, heuristic binary threshold models. This systemic approach successfully captured the broader regulatory architecture of Type 2 Diabetes, yielding a 133.3 percent increase in candidate target coverage and demonstrating that critical genetic risk is frequently mediated through peripheral systems, such as skeletal muscle and adipose tissue, rather than strictly through isolated pancreatic dysfunction.

Beyond generating a ranked portfolio of candidate effector genes, the rigorous validation protocols embedded within the pipeline ensure that the prioritized targets are biologically anchored and mathematically stable. The ability of the framework to successfully differentiate genuine metabolic regulatory signals from empirical non metabolic negative controls, coupled with high convergence on active epigenomic states and independent functional databases, provides a high degree of confidence in the top tier candidates, including JAZF1, FTO, and GRB14.

Ultimately, this probabilistic multi tissue integration bridges the critical gap between statistical genetic discovery and translational drug development. By mapping prioritized targets to systemic pleiotropic networks and evaluating their specific computational evidence for pharmacological tractability, the framework provides a pragmatic roadmap for experimental research. As multi omics repositories continue to expand, scalable and objective frameworks will be indispensable for generating high confidence, testable hypotheses that drive precision interventions for complex metabolic diseases.

## Statements and Declarations

### Data and Code Availability Statement

The datasets analyzed in this study were obtained from publicly accessible repositories and resources.

• Genome-Wide Association Study (GWAS) Resources:

NHGRI-EBI GWAS Catalog (https://www.ebi.ac.uk/gwas/)

• Expression Quantitative Trait Loci (eQTL) Resources:

Genotype-Tissue Expression (GTEx) Project v8 (https://gtexportal.org/)

• Epigenomic and Chromatin State Resources:

ENCODE Project (https://www.encodeproject.org/) and

Roadmap Epigenomics Project (https://egg2.wustl.edu/roadmap/web_portal/).

• Transcriptomic Validation Datasets:

Gene Expression Omnibus (GEO)

GSE38642 (https://www.ncbi.nlm.nih.gov/geo/query/acc.cgi?acc=GSE38642) and GSE81608 (https://www.ncbi.nlm.nih.gov/geo/query/acc.cgi?acc=GSE81608)

• External Validation Resources:

Open Targets Platform (https://platform.opentargets.org/)

• Pathway Enrichment Resources:

Enrichr (https://maayanlab.cloud/Enrichr/)

• Reference Panel:

1000 Genomes Project European reference panel (https://www.internationalgenome.org/).

All custom scripts, processed datasets, candidate gene rankings, pathway enrichment outputs, validation results, and figure-generation code used in this study are publicly available at: https://github.com/mohdmehboobuddin/CAS_X_T2D.

Due to repository size limitations, large raw datasets obtained from GTEx and other public resources are not hosted directly within the repository and should be downloaded from their original sources.

## CRediT Authorship Contribution Statement

**Mohd Mehboob Uddin:** Conceptualization, Methodology, Software, Data Curation, Formal Analysis, Investigation, Visualization, Validation, Writing – Original Draft, Writing – Review & Editing.

**Syed Mohd Zakariya Ali Khan:** Conceptualization, Methodology, Software, Data Curation, Formal Analysis, Investigation, Writing – Review & Editing.

**Bomma Yadagiri Kavitha:** Conceptualization, Methodology, Investigation, Visualization, Validation, Writing - Review & Editing.

## sDeclaration of Competing Interest

The authors declare that there are no known competing financial interests or personal relationships that could have appeared to influence the work reported in this study.

## Acknowledgements

The authors gratefully acknowledge the publicly available datasets and resources provided by the NHGRI-EBI GWAS Catalog, GTEx Consortium, Gene Expression Omnibus (GEO), Open Targets Platform, and Enrichr, which made this computational study possible. The authors extend their sincere gratitude to the principal of NIZAM College and also to the management of A.V. College of Arts, Science and Commerce, specifically the Correspondent, Dr. K. Gautami; the Principal, Dr. P. Padma; and the Director, M. Bhagvant Reddy, for their continuous institutional support and leadership.

## Ethical Approval

This study utilized only publicly available genomic, transcriptomic, and functional genomics datasets. No human participants, animal experiments, or identifiable personal data were directly involved in this research. Therefore, institutional ethical approval was not required.

