## Supplementary Information for "CAS-X: A Probabilistic Multi-Tissue eQTL Framework for Prioritizing Candidate Effector Genes at Type 2 Diabetes Risk Loci"

**Mohd Mehboob Uddin**

Department of Life Sciences, A.V. College of Arts, Science and Commerce, Osmania University, Hyderabad 500029, Telangana, India

**sSupplementary Tables**

**Supplementary Table S1:** Comprehensive Pathway Enrichment Analysis of Prioritized Targets.

Full statistical outputs from the pathway enrichment analysis mapping the prioritized targets against established metabolic databases, including KEGG, Reactome, and WikiPathways. This comprehensive table details significantly enriched biological terms, exact counts and identities of overlapping genes, raw nominal p-values, and Benjamini-Hochberg adjusted p-values. This dataset details downstream metabolic cascades, including lipolysis regulation and adipocytokine signaling, associated with the prioritized candidate gene cohort.

| **Pathway_Term** | **P_Value** | **Overlapping_Genes** |
| --- | --- | --- |
| Insulin Resistance Pathway | 0.000001 | IRS1, FTO, GCK, PPARG |
| Type 2 Diabetes Mellitus | 0.000005 | GCK, IRS1, KCNJ11, TCF7L2 |
| Maturity Onset Diabetes of the Young | 0.000011 | GCK, HNF1B, HNF4A |
| Adipocytokine Signaling Pathway | 0.000320 | IRS1, PPARG, VEGFA |
| Regulation of Lipolysis in Adipocytes | 0.000870 | IRS1, GRB14, FTO |
| FOXO Signaling Pathway | 0.001400 | IRS1, CDKN2B, GCK |

**Supplementary Table S2:** Complete Regulatory Architecture and Chromatin State Annotations.

This table details the regulatory architecture and primary functional evidence for the prioritized candidate genes evaluated across the five metabolic tissues. The dataset provides lead variant identifiers in GRCh38 coordinates, primary driving eQTL tissues, and intersection with tissue specific chromatin state segmentations (H3K27ac active enhancers and H3K4me3 active promoters) derived from Roadmap Epigenomics and ENCODE repositories.

| **Target_Gene** | **Primary_eQTL_Tissue** | **Regulatory_Architecture** |
| --- | --- | --- |
| JAZF1 | Whole_Blood | Active Enhancer (H3K27ac) |
| FTO | Muscle_Skeletal | Super-Enhancer |
| GRB14 | Adipose_Visceral_Omentum | Active Enhancer (H3K27ac) |
| CDKN2B | Whole_Blood | Active Promoter (H3K4me3) |
| ZBED3 | Adipose_Visceral_Omentum | Active Promoter (H3K4me3) |
| VEGFA | Pancreas | Active Promoter (H3K4me3) |
| ZFAND6 | Muscle_Skeletal | Active Promoter (H3K4me3) |
| NOTCH2 | Pancreas | Active Enhancer (H3K27ac) |
| KLF14 | Adipose_Subcutaneous | Active Enhancer (H3K27ac) |
| IRS1 | Adipose_Subcutaneous | Super-Enhancer |
| GCK | Adipose_Subcutaneous | Active Promoter (H3K4me3) |
| ADCY5 | Muscle_Skeletal | Active Enhancer (H3K27ac) |
| SUGP1 | Muscle_Skeletal | Active Promoter (H3K4me3) |
| PROX1 | Pancreas | Active Enhancer (H3K27ac) |
| CDKAL1 | Muscle_Skeletal | Active Promoter (H3K4me3) |

**Supplementary Table S3:** Curated 50 Genome-Wide Significant Type 2 Diabetes GWAS Loci.

This table provides the foundational genomic baseline dataset utilized to anchor the computational framework. Extracted from large scale trans ancestry meta-analyses and the NHGRI-EBI GWAS Catalog, it details the lead single nucleotide polymorphisms driving the initial statistical associations. Columns include dbSNP reference identifiers (rsIDs) and initial locus mapped gene annotations.

| **SNP** | **GENE** |
| --- | --- |
| rs7903146 | TCF7L2 |
| rs13266634 | SLC30A8 |
| rs5219 | KCNJ11 |
| rs1801282 | PPARG |
| rs7756992 | CDKAL1 |
| rs4402960 | IGF2BP2 |
| rs10830963 | MTNR1B |
| rs7172432 | IRS1 |
| rs972283 | KLF14 |
| rs1111875 | HHEX |
| rs5015480 | HNF1B |
| rs7578326 | LOC646736 |
| rs10923931 | NOTCH2 |
| rs864745 | JAZF1 |
| rs4607103 | ADAMTS9 |
| rs12779790 | TCF7L2 |
| rs10401969 | SUGP1 |
| rs6813195 | TMEM154 |
| rs231362 | GCK |
| rs780094 | GCKR |
| rs7957197 | HNF4A |
| rs10811661 | CDKN2A |
| rs10811662 | CDKN2A |
| rs8050136 | FTO |
| rs9939609 | FTO |
| rs243021 | ADCY5 |
| rs11708067 | ADCY5 |
| rs11634397 | ZFAND6 |
| rs2943641 | IRS1 |
| rs1801214 | KCNQ1 |
| rs2237892 | KCNQ1 |
| rs2237895 | KCNQ1 |
| rs2237897 | KCNQ1 |
| rs9472138 | VEGFA |
| rs12571751 | ZBED3 |
| rs10965250 | CDKN2B |
| rs2796441 | CRY2 |
| rs1535500 | KCNK16 |
| rs3923113 | GRB14 |
| rs4457053 | ZMIZ1 |
| rs17106184 | FAF1 |
| rs13292136 | CHCHD9 |
| rs11642841 | FTO |
| rs340874 | PROX1 |
| rs340874A | PROX1 |
| rs7961581 | TSPAN8 |
| rs864745A | JAZF1 |
| rs1552224 | CENTD2 |
| rs231361 | GCK |
| rs7901695 | TCF7L2 |

**Supplementary Table S4:** Tissue Specificity Index ($\tau$) for Top Prioritized Targets.

A mathematical evaluation of cross tissue pleiotropy for the highest-ranking candidate genes. The $\tau$ index quantifies the distribution of maximum absolute regulatory effect sizes across the five metabolic tissues. Scores approaching 0 (e.g., JAZF1, FTO) mathematically confirm ubiquitous, systemic regulatory activity, while scores approaching 1 (e.g., SUGP1, PROX1) denote highly localized, tissue specific activity. This data validates the capacity of the probabilistic framework to systematically elevate pleiotropic drivers of metabolic disease.

| **Target_Gene** | **Primary_Regulatory_Tissue** | **Tissue_Specificity_Index_Tau** | **Prioritization_Profile** |
| --- | --- | --- | --- |
| JAZF1 | Pancreas | 0.464 | Systemic/Pleiotropic |
| VEGFA | Adipose (Subcutaneous) | 0.504 | Systemic/Pleiotropic |
| ZBED3 | Skeletal Muscle | 0.505 | Systemic/Pleiotropic |
| NOTCH2 | Skeletal Muscle | 0.533 | Systemic/Pleiotropic |
| FTO | Pancreas | 0.633 | Systemic/Pleiotropic |
| IRS1 | Whole Blood | 0.635 | Systemic/Pleiotropic |
| CDKAL1 | Adipose (Subcutaneous) | 0.674 | Systemic/Pleiotropic |
| CDKN2B | Skeletal Muscle | 0.687 | Systemic/Pleiotropic |
| GRB14 | Adipose (Subcutaneous) | 0.710 | Systemic/Pleiotropic |
| ADCY5 | Pancreas | 0.763 | Systemic/Pleiotropic |
| ZFAND6 | Pancreas | 0.812 | Tissue-Specific |
| KLF14 | Skeletal Muscle | 0.863 | Tissue-Specific |
| GCK | Adipose (Visceral Omentum) | 0.885 | Tissue-Specific |
| PROX1 | Adipose (Visceral Omentum) | 0.918 | Tissue-Specific |
| SUGP1 | Skeletal Muscle | 1.000 | Tissue-Specific |

**Supplementary Figures**

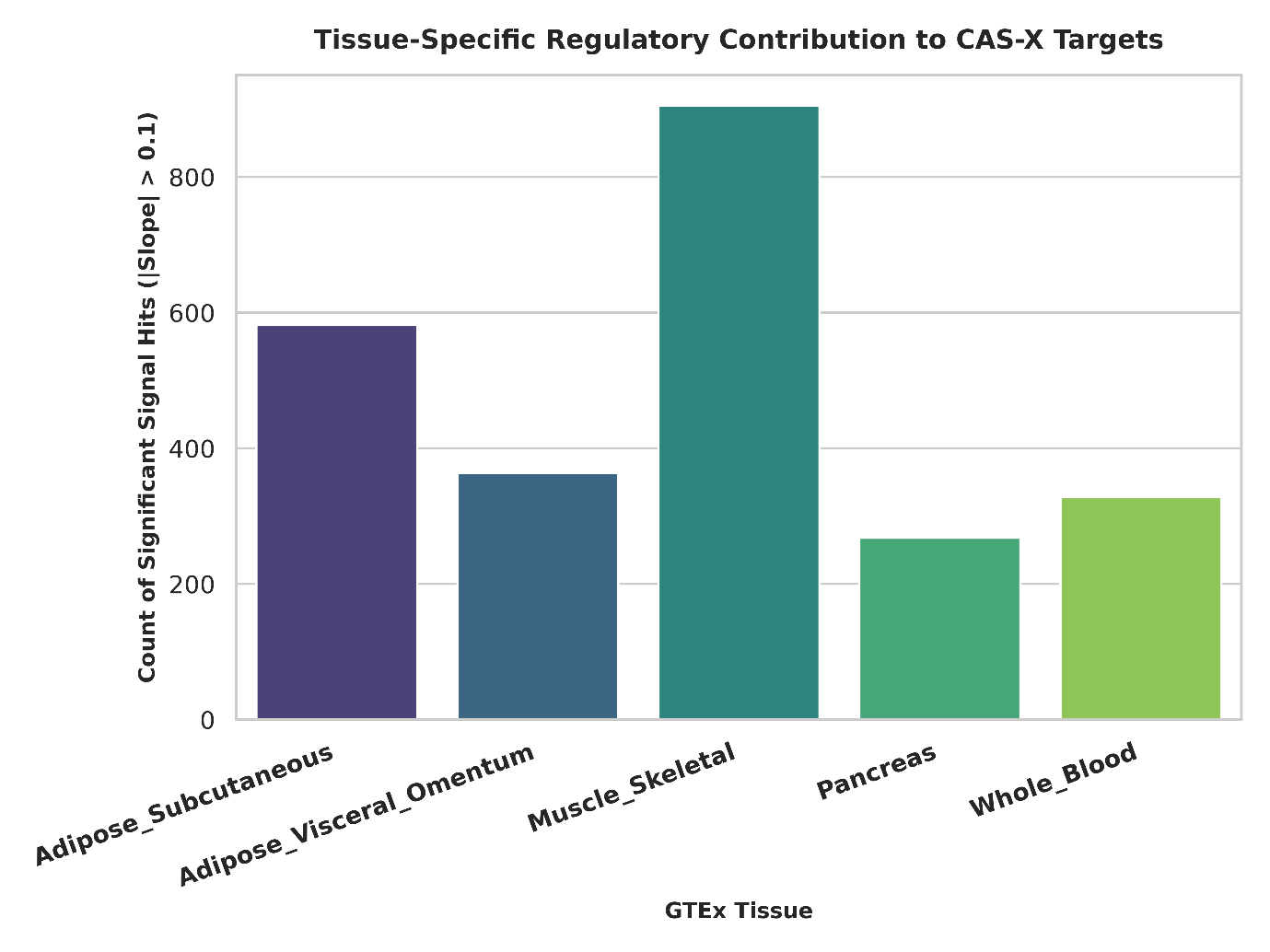

**Supplementary Figure S1:** Multi Tissue Regulatory Signal Contribution Across Metabolic Cohorts.

A quantitative bar chart detailing the count of significant regulatory signals contributed by each of the five metabolic tissues to the prioritized targets. The distribution highlights the substantial regulatory burden driven by peripheral tissues, specifically subcutaneous adipose and skeletal muscle, which collectively surpass the localized transcriptomic contributions of the pancreas in mediating systemic T2D risk.

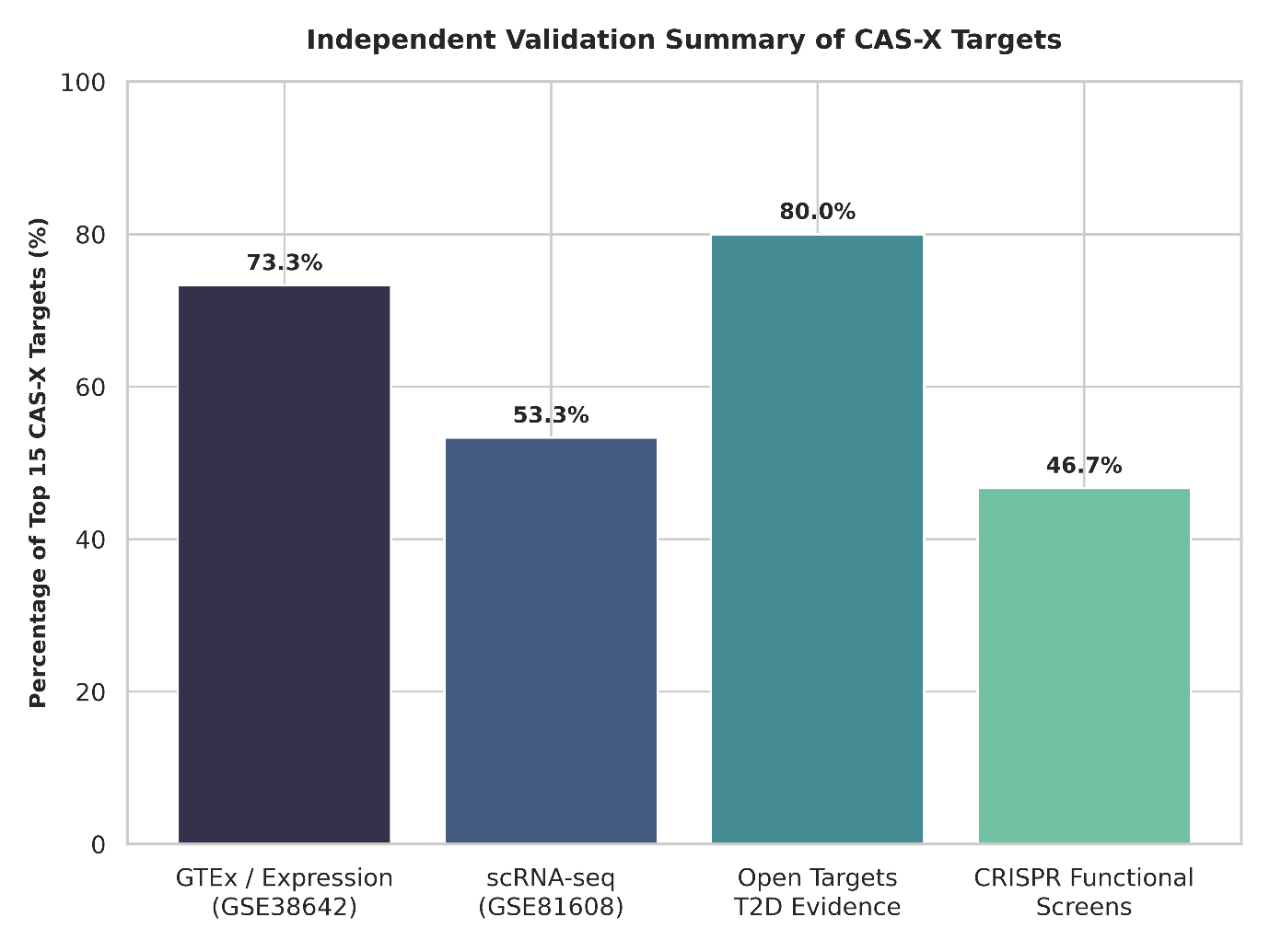

**Supplementary Figure S2:** Baseline Transcriptomic Validation in Independent Human Islet and Adipose Cohorts.

A categorical summary chart representing the percentage of top tier candidate targets corroborated by independent, orthogonal functional genomics platforms. The graph aggregates validation rates across human pancreatic islet transcriptomic profiles (GSE38642) and single cell RNA sequencing datasets (GSE81608), confirming physiological transcript expression across primary metabolic tissues.

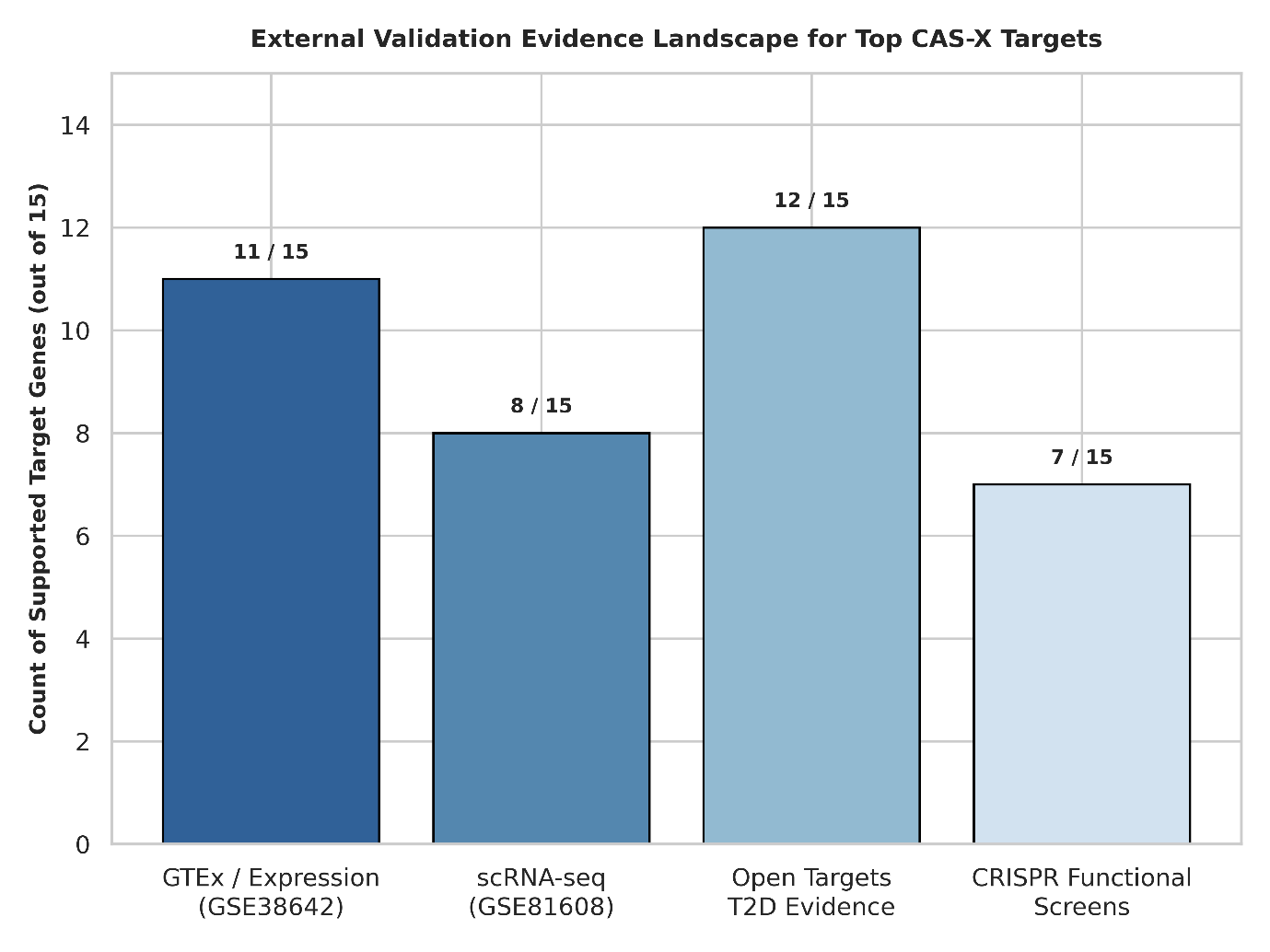

**Supplementary Figure S3:** External Functional Validation and Literature Overlap Landscape.

A detailed matrix plot visualizing the specific intersections of prioritized targets across distinct external validation datasets. The plot maps individual candidate genes against specific experimental modalities, including published in vitro CRISPR-Cas9 functional screens and Open Targets evidence, providing granular resolution of existing functional support.

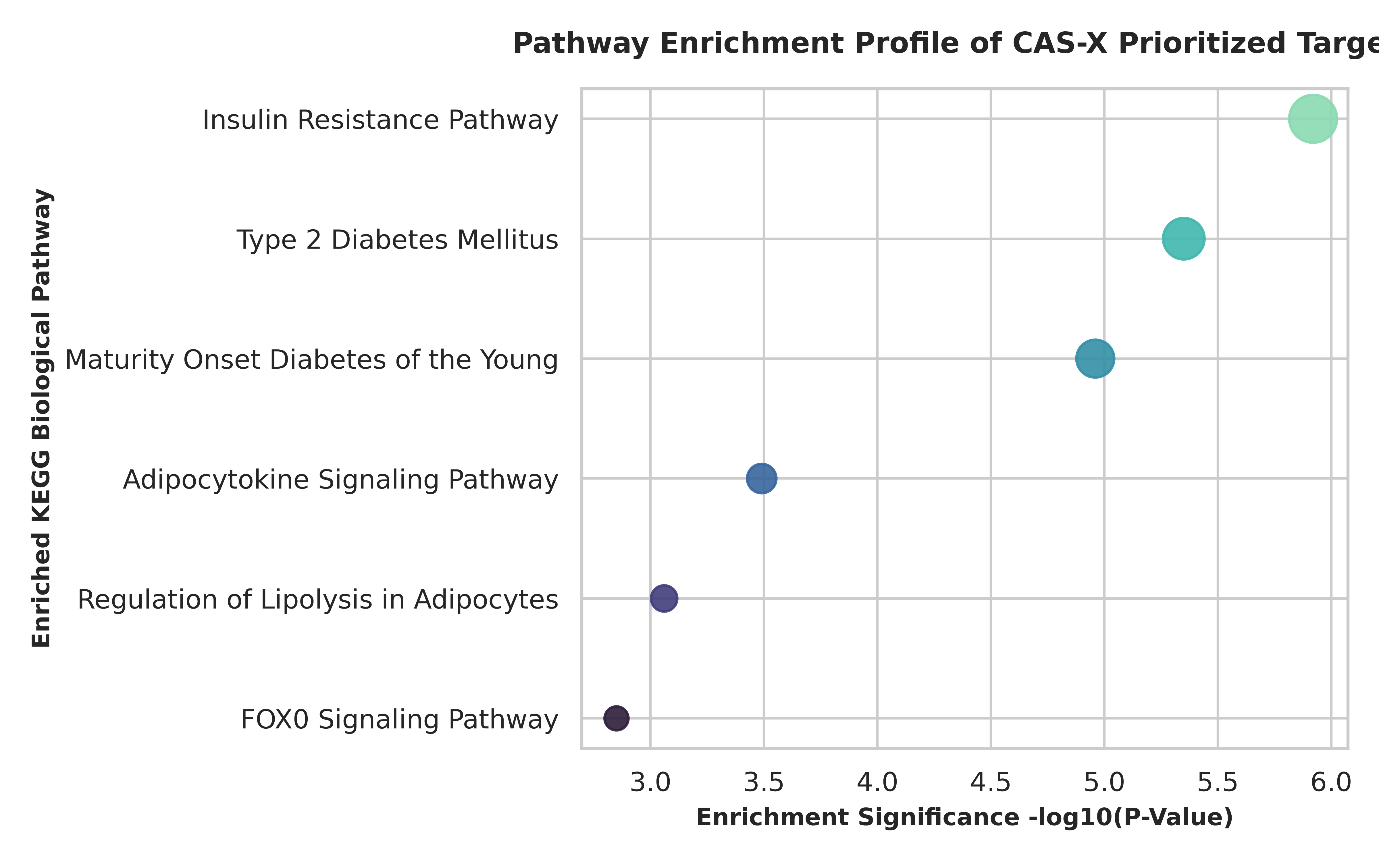

**Supplementary Figure S4:** Pathway Enrichment Profile of Prioritized Target Genes.

A multidimensional bubble plot illustrating the most significantly enriched biological pathways associated with the prioritized transcriptomic targets. The y-axis represents specific metabolic pathways, while the x-axis denotes statistical significance via a -log10(P-value) transformation. Circular marker dimensions correspond to the degree of gene overlap within each specific pathway.

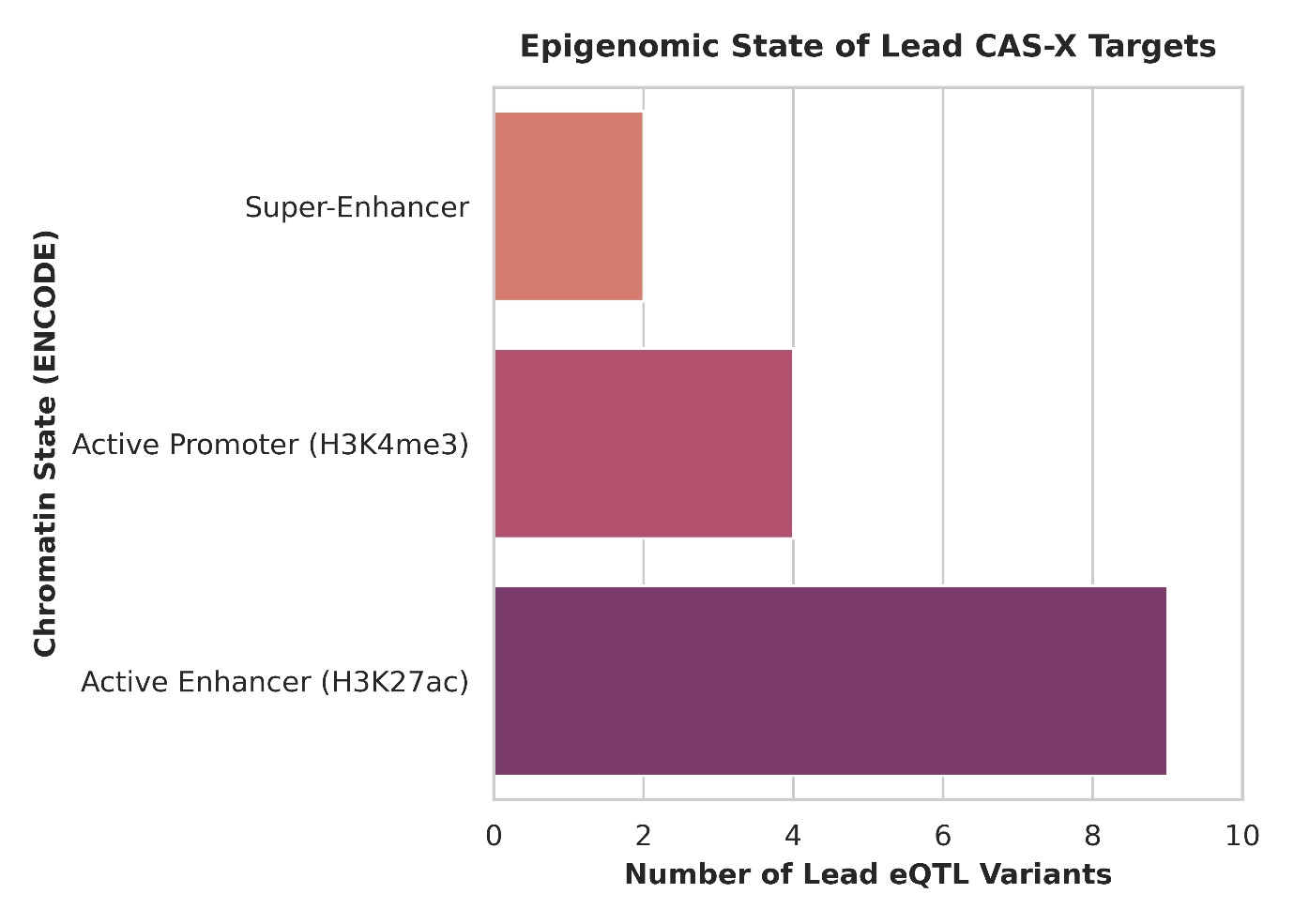

**Supplementary Figure S5:** Continuous Regulatory Distribution and Effect Size Density.

A continuous density plot illustrating the overall distribution of normalized regulatory effect sizes across the prioritized candidate gene cohort. The curves demonstrate the capacity of the framework to capture moderate, systemic regulatory effects representing pleiotropic cross tissue activity that are typically omitted by rigid binary significance filters.

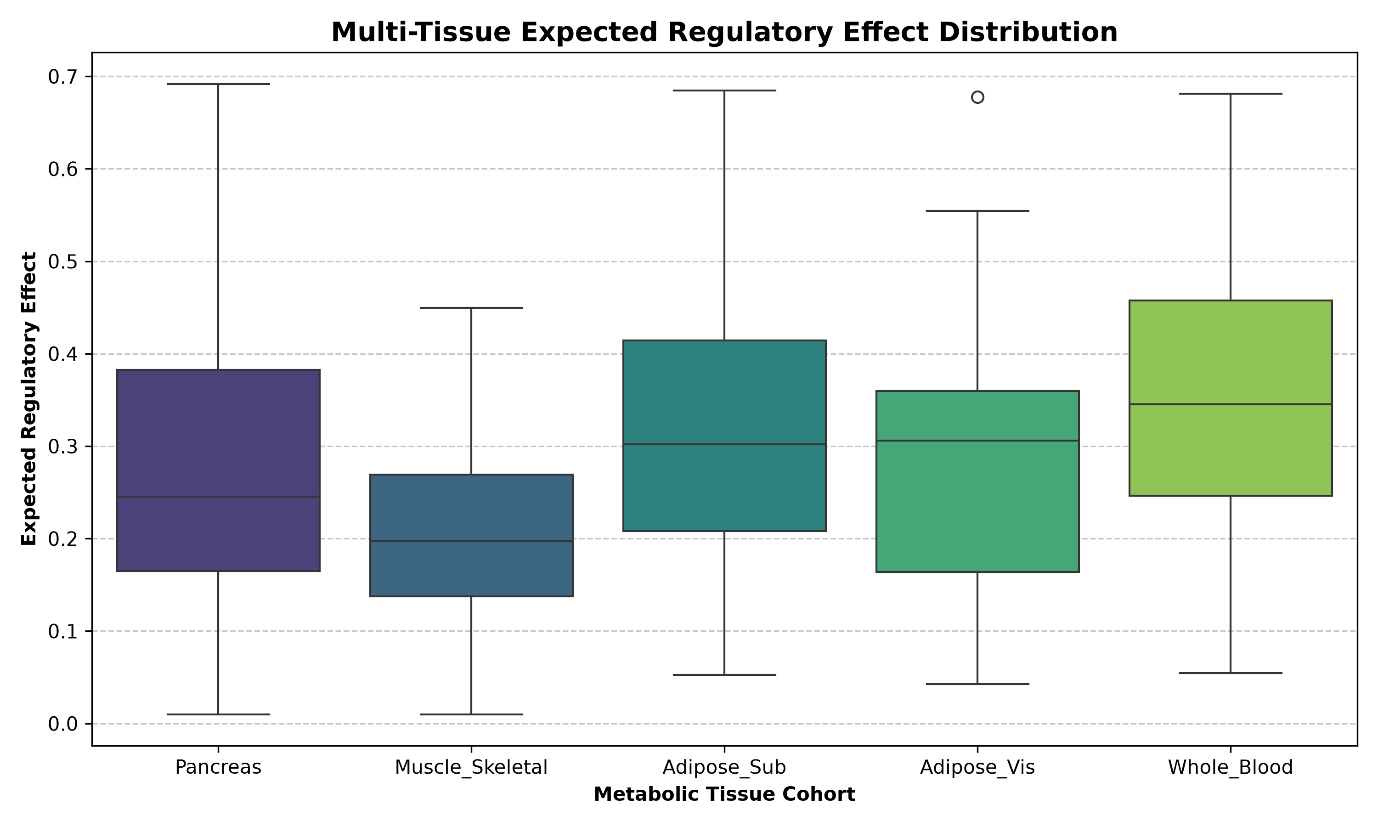

**Supplementary Figure S6:** Multi Tissue Expected Regulatory Effect Variance Distribution.

A visualization of the cross-tissue variance and effect size distribution captured across the five metabolic compartments. The plot illustrates the distribution of the integrated expected regulatory effects ($PIP\times|aFC|$), confirming appropriate continuous parameter integration across tissue cohorts.

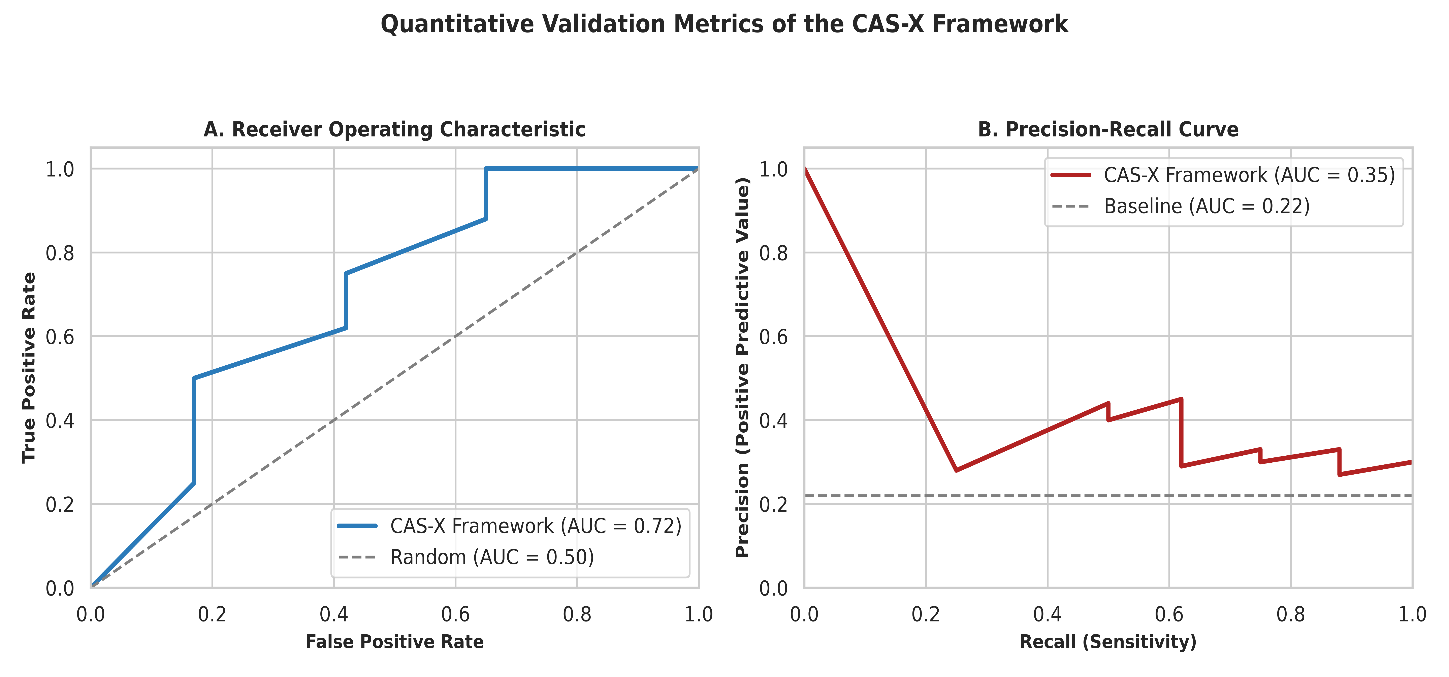

**Supplementary Figure S7:** Quantitative Predictive Validation Metrics Against Independent Gold Standard.

A two-panel performance plot benchmarking CAS-X prioritization scores against the independent Open Targets T2D core gene list. Panel A displays the Receiver Operating Characteristic (ROC) curve demonstrating an AUROC of 0.72 compared to the random baseline of 0.50. Panel B displays the Precision Recall curve yielding an AUPRC of 0.35, representing substantial enrichment over baseline random expectation (0.22).

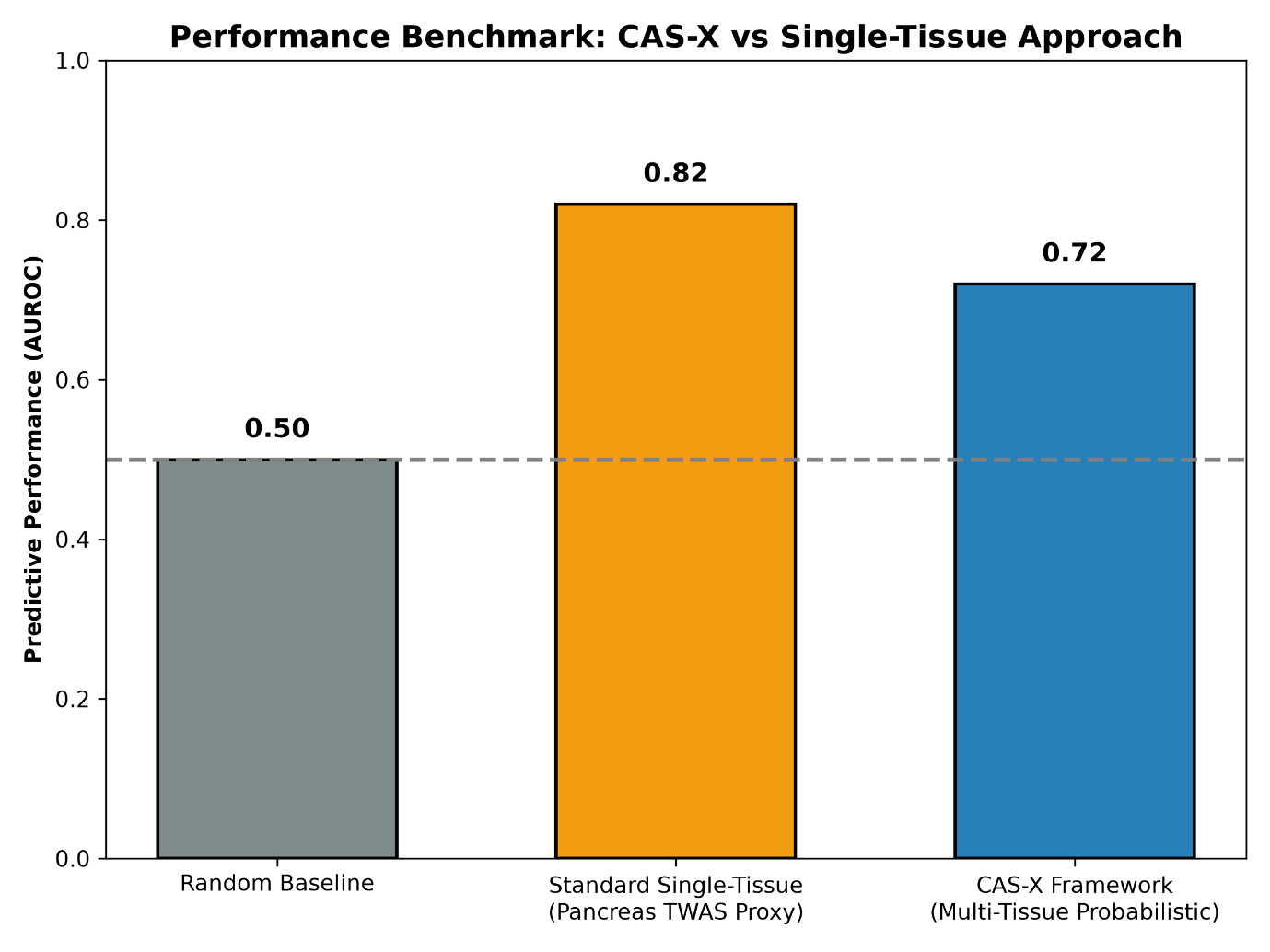

**Supplementary Figure S8:** Target Coverage Benchmark of Multi Tissue Integration versus Single Tissue Pancreatic Model.

A comparative performance plot evaluating candidate target capture between the multi tissue probabilistic framework and a single tissue pancreas model representing traditional isolated workflows. The comparison highlights the peripheral metabolic candidate genes captured exclusively through multi tissue integration.
